# A *Nanog*-dependent gene cluster initiates the specification of the pluripotent epiblast

**DOI:** 10.1101/707679

**Authors:** Nicolas Allègre, Sabine Chauveau, Cynthia Dennis, Yoan Renaud, Lorena Valverde Estrella, Pierre Pouchin, Michel Cohen-Tannoudji, Claire Chazaud

## Abstract

The epiblast (Epi) is the source of embryonic stem (ES) cells and all embryonic tissues. It differentiates alongside the primitive endoderm (PrE) in a randomly distributed “salt and pepper” pattern from the inner cell mass (ICM) during preimplantation of the mammalian embryo. NANOG and GATA6 are key regulators of this binary differentiation event, which is further modulated by heterogeneous FGF signalling. When and how Epi and PrE lineage specification is initiated within the developing embryo is still unclear. Here we generated NANOG and GATA6 double KO (*DKO*) mouse embryos and performed single-cell expression analyses. We found that the ICM was unable to differentiate in the *DKO* mice, allowing us to characterize the ICM precursor state. The normally heterogeneous expression of *Fgf4* between cells was significantly reduced in *DKO* ICMs, impairing FGF signalling. In contrast, several pluripotency markers did still display cell-to-cell expression variability in *DKO* ICMs. This revealed a primary heterogeneity independent of NANOG, GATA6 and FGF signalling that may also be conserved in humans. We found that NANOG is key in the initiation of epiblast specification already between the 16- and 32-cell stages, enabling the cell-clustered expression of many pluripotency genes. Our data uncover previously unknown biology in the early mouse embryo with potential implications for the field of pluripotent stem cells in human and other mammals.

---

During preimplantation, the mammalian embryo develops into a blastocyst via two cell lineage differentiation events. First, from the 16-cell (16C) stage, inner cells segregate from outer cells and form the inner-cell mass (ICM), while the outer cells produce the extraembryonic trophectoderm (TE). A second differentiation event occurs within the ICM, with the production of the epiblast (Epi), the source of embryonic stem (ES) cells and of all embryonic tissues, and primitive endoderm (PrE) cells. At the beginning of blastocyst formation, around the 20C-32C stage, all ICM cells coexpress NANOG and GATA6. Subsequently, between the 32C and 90C stage, these cells asynchronously differentiate into Epi, expressing solely NANOG, or PrE expressing only GATA6, in an apparently random pattern (Chazaud et al., 2006; Guo et al., 2010; Ohnishi et al., 2014; Plusa et al., 2008; Saiz et al., 2016). As single *Gata6* and *Nanog* mutations prevent the specification of PrE and Epi respectively (Bessonnard et al., 2014; Frankenberg et al., 2011; Schrode et al., 2014), we hypothesized that in the absence of both *Nanog* and *Gata6*, ICM cells should not be able to differentiate and should remain in a precursor state, with or without cell heterogeneity. We thus generated *N*^-/-^;*G*^-/-^ double KO (*DKO*) embryos for analysis. After validating that TE and ICM were properly segregated (Supp data Fig.1a), we examined whether ICM cells could still differentiate in these compound mutant embryos by immunofluorescence (IF) analysis of PrE or Epi specific genes. We also performed RTqPCR on whole individual ICMs from wild type (*WT*), single KOs (*N*^-/-^ and *G*^-/-^) and *DKO* embryos at the 32-cell stage (32C), when some ICM cells normally start to differentiate, and at stage 90C when most of the cells have acquired either an Epi or a PrE identity (Fig. 1a) (Bessonnard et al., 2014; Plusa et al., 2008; Saiz et al., 2016). IF (Supp data Fig.1a,b) and RTqPCR (Fig. 1b; supplemental information (SI) 1) showed that none of the eight PrE markers tested (e.g., *Sox17, Pdgfra, Gata4, Foxq1…*) (Artus et al., 2011; Ohnishi et al., 2014; Plusa et al., 2008) were expressed in the double mutants. Thus, PrE does not differentiate in these embryos. With the 16 Epi markers, RTqPCR revealed different expression patterns (Fig. 1b; SI 1): expression of some of the markers (e.g., *Nanog, Bmp4, Tdgf1*) was significantly down regulated or absent in *DKO* ICMs from either 32C or 90C stages, while others remained constant (*Zfp42, Prdm14*) or increased (*Klf2*). The absence of *Nanog* RNA in *DKO* embryos indicated that NANOG is required for its own expression in embryos, which is different than the self-repression reported in ES cells (Fidalgo et al., 2012; Navarro et al., 2012). Altogether, our gene expression analyses revealed that *DKO* ICM cells cannot differentiate into Epi or PrE.

**Figure 1:**
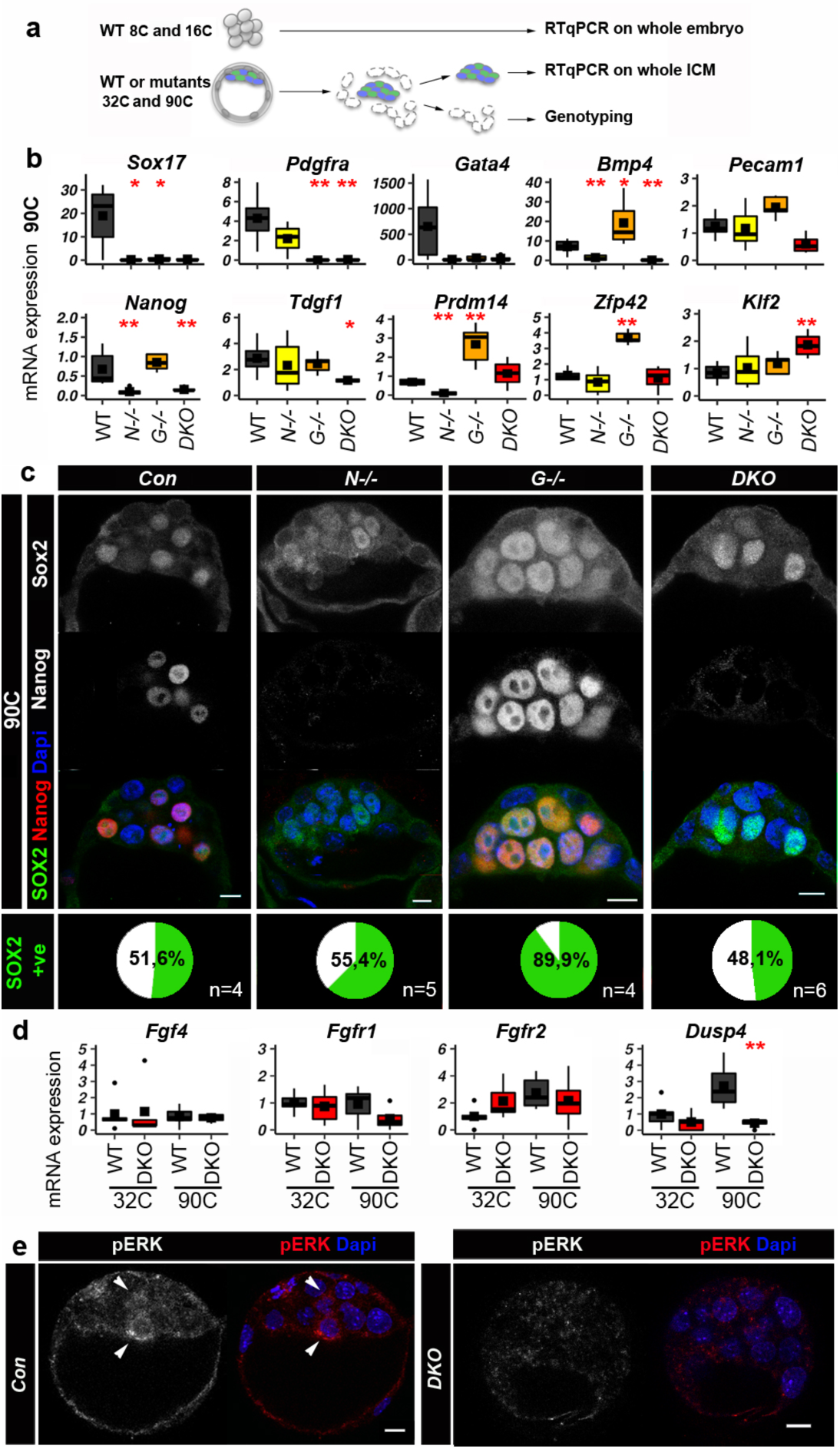
Absence of PrE and Epi differentiation in *DKO* embryos. **a**, Schematic representation of the experimental procedure used in **b, d** and SI 1. For 8C and 16C stages, individual whole embryos were analysed whereas for 32C and 90C stages individual whole ICM were isolated by immunosurgery. **b**, Expression analysis of some PrE and Epi genes by RTqPCR. Boxplots showing the relative RNA levels in 90C *WT* (n=6), *N*^-/-^ (n=6), *G*^-/-^ (n=5) and *DKO* (n= 4) whole ICMs. The mean is represented by a square, the median by the central line. The levels are relative to the 32C *WT* mean (see SI 1). Red asterisks indicate the statistical significance compared to the *WT* sample: *p<0,05, **p<0,01 (Wilcoxon test, see SI 8 for all values). **c**, Representative immunofluorescence confocal images of single mutant, *DKO* and control (*Con*) embryos at 90C stage, showing the localisation of NANOG and SOX2. Percentages of SOX2 expressing cells (+ve) in the ICM are indicated on the bottom panel. Scale bars: 10 µm. **d**, RTqPCR analysis of some FGF pathway members on *WT* and *DKO* ICMs at 32C (*WT* n=5, *N*^-/-^ n=5, *G*^-/-^ n= 8, *DKO* n=5) and 90C stages as in (**b** and SI 1). **e**, Representative immunofluorescence confocal images of *DKO* (n=5) and control embryos at stage 64-90C to show pERK presence (arrowheads). Scale bars: 10 µm.

To characterize the undifferentiated state of *DKO* ICM cells in more detail we compared the expression patterns of various markers with those in the *WT* embryos at different stages. We first examined expression of the pluripotency markers SOX2 and PECAM1, as they are present at stage 16C in inner cells only, during formation of the ICM, before being progressively restricted to Epi cells within the ICM at the 90C stage(Avilion et al., 2003; Fiorenzano et al., 2016; Robson et al., 2001; White et al., 2016; Wicklow et al., 2014). Both SOX2 and PECAM1 were expressed in *DKO* embryos at 32C (n=4) and 90C (n=7) stages (Figure 1c, Supp data Fig.1d), with SOX2 displaying the same heterogeneous nuclear levels in the characteristic salt and pepper pattern as in *WT* embryos. This heterogeneous pattern was completely lost in the *G*^-/-^ embryos, where all cells of the ICM differentiate into Epi cells. This indicates that at this later stage, heterogeneous SOX2 expression is influenced by the universal presence of NANOG and/or loss of GATA6, whereas at the 16C stage, before the ratio is established, it must be controlled differently. Thus, SOX2 expression is controlled by different inputs that are first independent and then dependent on a high NANOG/GATA6 expression ratio. SOX21 is another early marker of pluripotency that is heterogeneously expressed at the 4C stage(Goolam et al., 2016). At the 90C stage, we found that SOX21 levels decreased in *WT* ICM cells compared to earlier stages(Goolam et al., 2016) and indeed this protein in undetectable in ESCs(Moretto Zita et al., 2015) that correspond to late blastocyst Epi cells(Boroviak et al., 2014). Interestingly, *DKO* ICM cells exhibited a heterogeneous pattern of SOX21 (Supp data Fig.1c), indicating the maintenance of the early expression pattern. Altogether, our results show that in the absence of NANOG and GATA6, the ICM remains at a precursor state that still displays cell heterogeneity, illustrated by the expression patterns of SOX2 and SOX21.

We hypothesized that this ICM heterogeneity in *DKO* mutants may be due to FGF signalling. *Fgf4* is specifically expressed in Epi cells (Frankenberg et al., 2011; Kurimoto et al., 2006), and is one of the earliest genes that shows binary expression among ICM cells, around the 32C stage (Guo et al., 2010; Ohnishi et al., 2014). *Fgf4* expression is not detected in *N*^-/-^ embryos (Frankenberg et al., 2011), due either to the lack of activation by NANOG, or to repression by GATA6, or both. Indeed, these two transcription factors directly bind the *Fgf4* locus, respectively in undifferentiated (Chen et al., 2008; Gagliardi et al., 2013) or PrE differentiated (Wamaitha et al., 2015) ES cells. Thus, after the 32C stage, *Fgf4* heterogeneous expression is maintained by a high NANOG/GATA6 ratio in Epi cells. However, it is unknown whether *Fgf4* expression is solely induced by this high ratio or is initially independent from it, and could thus be driving the ICM heterogeneity seen in the *DKO* mutants. In *WT* embryos, very low levels of *Fgf4* transcripts were detected at 8C and 16C, while increased expression was observed in 32C and 90C ICMs (SI 1). In *G*^-/-^ mutants, *Fgf4* expression was further induced, while it decreased in *N*^-/-^ embryos (SI 1). This is similar to ES cells where *Nanog* is required for producing high levels of *Fgf4* transcripts (Festuccia et al., 2012). In contrast to the single KOs, the level of *Fgf4* expression in the *DKO* mutants was similar to *WT*’s at the 32C and 90C stages (Fig.1d). This implies that *Nanog* is not required for *Fgf4* expression in early ICM cells. Moreover, *Fgfr1* and *Fgfr2* RNAs were also present in the double mutants, enabling FGF pathway signalling (Fig. 1d). However, in contrast to the *WT*, expression of FGF signalling target genes such as *Dusp4, Etv4* and *Spry4* (Kang et al., 2017; Morgani et al., 2018) were down-regulated in *DKO* mutants at the 90C stage (Fig. 1d, Supp data Fig.1e, SI 1). Thus, when analysing the entire *DKO* ICM cell population, FGF signalling is not activated, despite *Fgf4* and *Fgfrs* expression levels similar to those seen in the *WT* ICM, where FGF signalling is activated. Accordingly, ERK phophorylation that has been described in some precursor and PrE cells (Azami et al, *Development*, in press) of *WT* embryos was also absent in *DKO* ICMs (Fig. 1e, n= 5). A possible explanation for this is that, in contrast to *WT* embryos that display heterogeneous *Fgf4* expression across ICM cells (Guo et al., 2010; Ohnishi et al., 2014), *DKO* ICM cells may all express similar levels of *Fgf4*, which would then be comparatively lower than in the *WT* cells. In *WT* ICMs, the differential expression would locally create high levels that reach the threshold required for inducing FGF signalling. In contrast, a globally similar but homogeneous level of FGF4 throughout the ICM may be locally lower in *DKO* embryos, and possibly insufficient for downstream pathway activation. To address this further, we carried out single cell gene expression analyses.

Before analysing mutant cells, we performed a quantitative analysis of gene expression in single cells of *WT* embryos as a reference (*WT*-Ref) at different time points of development (16C, 32C, 64C and 90C stages) by single-cell RNA analyses (Fig. 2a,b). We focused our study on known markers for PrE and Epi (Chazaud et al., 2006; Fiorenzano et al., 2016; Guo et al., 2010; Kurimoto et al., 2006; Ohnishi et al., 2014; Plusa et al., 2008) genes, reported to be expressed before the 32C stage and/or linked to pluripotency in embryos or ES cells (e.g., *Zscan4, Tcfcp2l1, Sox21, Prdm14*) (Burton et al., 2013; Falco et al., 2007; Goolam et al., 2016; Pelton et al., 2002), and FGF pathway genes (Kang et al., 2017; Ohnishi et al., 2014). Unsupervised hierarchical clustering showed that the cells regrouped according to their stage and level of differentiation (Supp data Fig. 2a). The progressive evolution of differentiation toward Epi and PrE states was also captured by principal component analysis (PCA) (Fig. 2c). Developmental time segregated cells along PC1, whereas PC2 highlighted differentiation between Epi and PrE states, which can be depicted by the levels of *Fgf4* expression (Fig. 2d), and by several Epi and PrE markers (Supp data Fig. 2b,2c). We then assessed the dynamics of expression for each gene at each stage (Supp data Fig. 2c, SI 2). Except for *Gata6* and *Fgfr2*, the analysed PrE markers initiated their expression rather late between the 64C and 90C stage, consistent with their induction by a low NANOG/GATA6 ratio (Bessonnard et al., 2014; Ohnishi et al., 2014; Saiz et al., 2016). Several Epi/pluripotency markers such as *Fgf4, Prdm14, Klf2, Klf4, Tdgf1* displayed heterogeneous expression among ICM cells at the 32C stage (SI 2). As previously published (Guo et al., 2010; Ohnishi et al., 2014), our results showed that *Fgf4* expression was low at the 16C and increased from the 32C stage in a subset of cells (SI 2), segregating the samples into two populations of *Fgf4*-expressing (*Fgf4+*ve/Epi) and non-expressing (*Fgf4–*ve/PrE) cells (Fig. 2e). Consequently, for each stage we subdivided the cells into *Fgf4*+ve and *Fgf4–*ve expression subgroups (Fig. 2f; SI 3). This enabled us to visualise differences between Epi and PrE cells at the 64C and 90C stage, in agreement with previous reports (Guo et al., 2010; Kang et al., 2017; Ohnishi et al., 2014). At the 32C stage, *Prdm14, Klf2, Klf4, Tdgf1, Sox21, Sox2* and *Nanog* already displayed differential expression between *Fgf4*+ve and *Fgf4–*ve cells, whereas other Epi markers like *Bmp4, Zfp42, Enox1* or *Esrrb* exhibited a later restriction to the Epi lineage (Fig. 2f and SI 3). We also examined the expression of some components of the FGF pathway (SI 3). As previously described (Kang et al., 2017; Molotkov et al., 2017; Ohnishi et al., 2014) *Fgfr1* was expressed at similar levels between Epi (*Fgf4*+ve) and PrE (*Fgf4*–ve) cells, while *Fgfr2* expression became restricted to PrE cells rather late, around the 90C stage. This confirms that all ICM cells of the early blastocyst (∼32C) are equally able to receive the FGF signal through FGFR1 (Kang et al., 2017; Molotkov et al., 2017) and that the beginning of ICM cell differentiation most likely relies only on the local FGF4 concentration.

**Figure 2:**
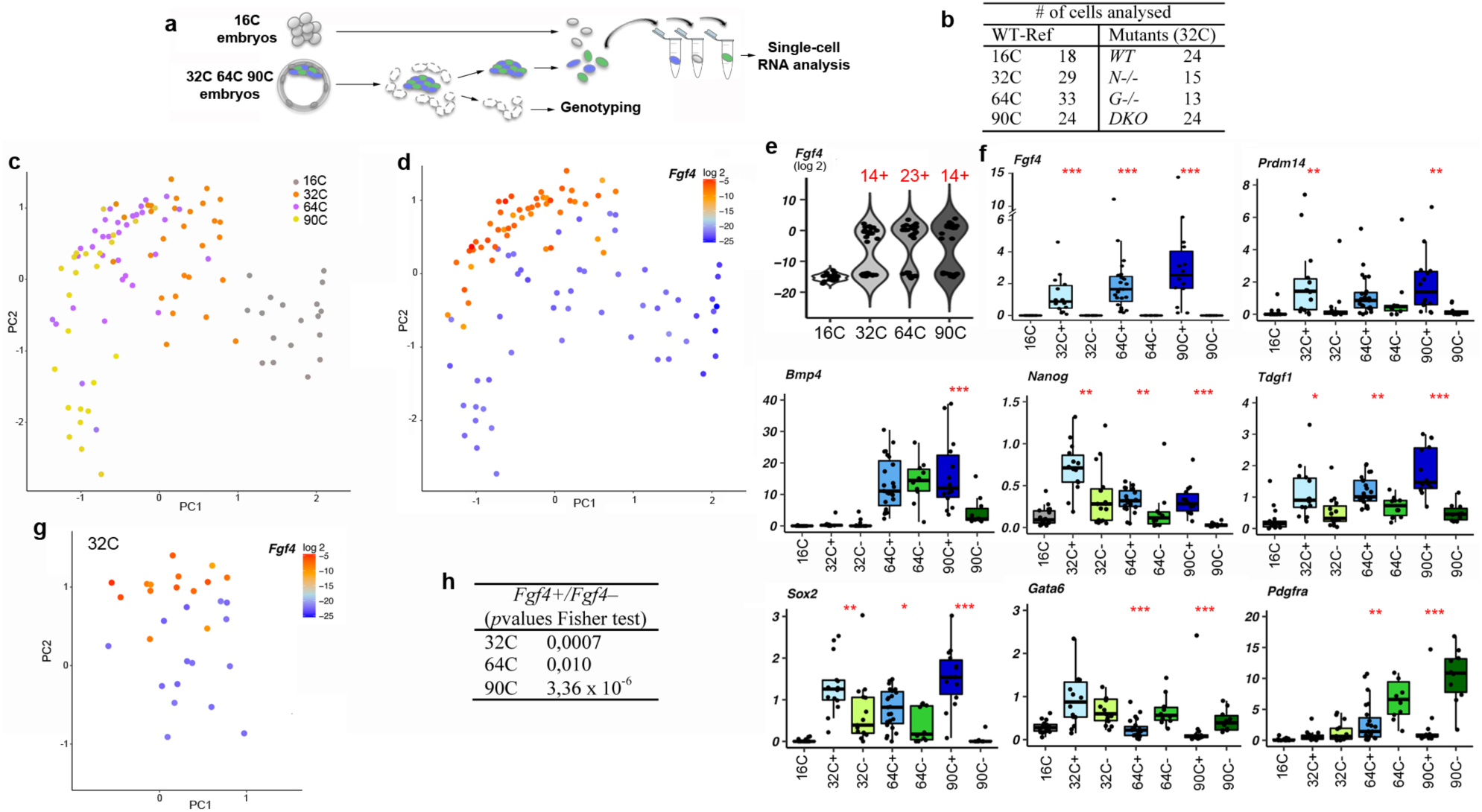
Single-cell RNA analysis reveals the emergence of two cell populations at stage 32C in *WT* embryos. **a**, Procedure for single cell isolation. **b**, Number of cells analysed in the CD1 (*WT*-Ref) and transgenic backgrounds. **c**, PCA map where stages are represented by different colours (PC1 score= 24,40%, PC2 score = 14,77%). **d**, PCA plot shown in (**c**) with graded colours indicating the level of *Fgf4* expression in each cell at the four stages. **e**, Violin plot representation of expression of *Fgf4* in individual cells. The number of *Fgf4*+ve cells are indicated in red. **f**, Boxplots showing the single-cell expression levels (fold change relative to the mean of 32C stage cells) of indicated genes at the four different stages, subdivided in *Fgf4*+ve and *Fgf4*–ve subpopulations, according to the violin plots shown in **e**. Red asterisks indicate statistically significant differences between *Fgf4*+ve and *Fgf4*–ve cells: *p<0,05, **p<0,01, ***p<0,001 (Wilcoxon test, see SI 8 for all values). **g**, Same PCA plot as in (**d**), showing only 32C stage cells. **h**, Fisher’s exact test for *Fgf4*+ve and *Fgf4*–ve cells distribution (**c**) after repeated k-means clustering (k=2) (**g**).

Strikingly, *Fgf4+ve* cells were already significantly clustered at stage 32C on the PCA map (Fig. 2d, 2g, 2h). This indicated that *Fgf4* expression is not stochastic at this stage and that expression of many other genes participate to segregate *Fgf4* expressing cells on the map. Indeed, several Epi/pluripotency genes displayed a similar distribution (Supp data Fig. 2c, 3b) and the *Fgf4+*ve cells on the PCA map were mainly included within *Nanog*-, *Prdm14-, Klf2-, Klf4-* or *Tdgf1*-expressing populations (Supp data Fig. 3a, 3d). Accordingly, significant correlations of expression were found between *Fgf4* and 8 Epi/pluripotency genes at 32C (Supp data Fig. 3c). Thus, *Fgf4* expression is possibly induced by the combined activity of some of these factors, or by other unknown factor(s), restricting the expression of these genes to a subset of cells. Altogether, these results show that already at the 32C stage, a group of cells share a gene expression signature comprising several Epi/pluripotency markers together with *Fgf4*. This strongly suggests that the Epi state emerges earlier than previously thought, between the 16C and 32C stages.

**Figure 3:**
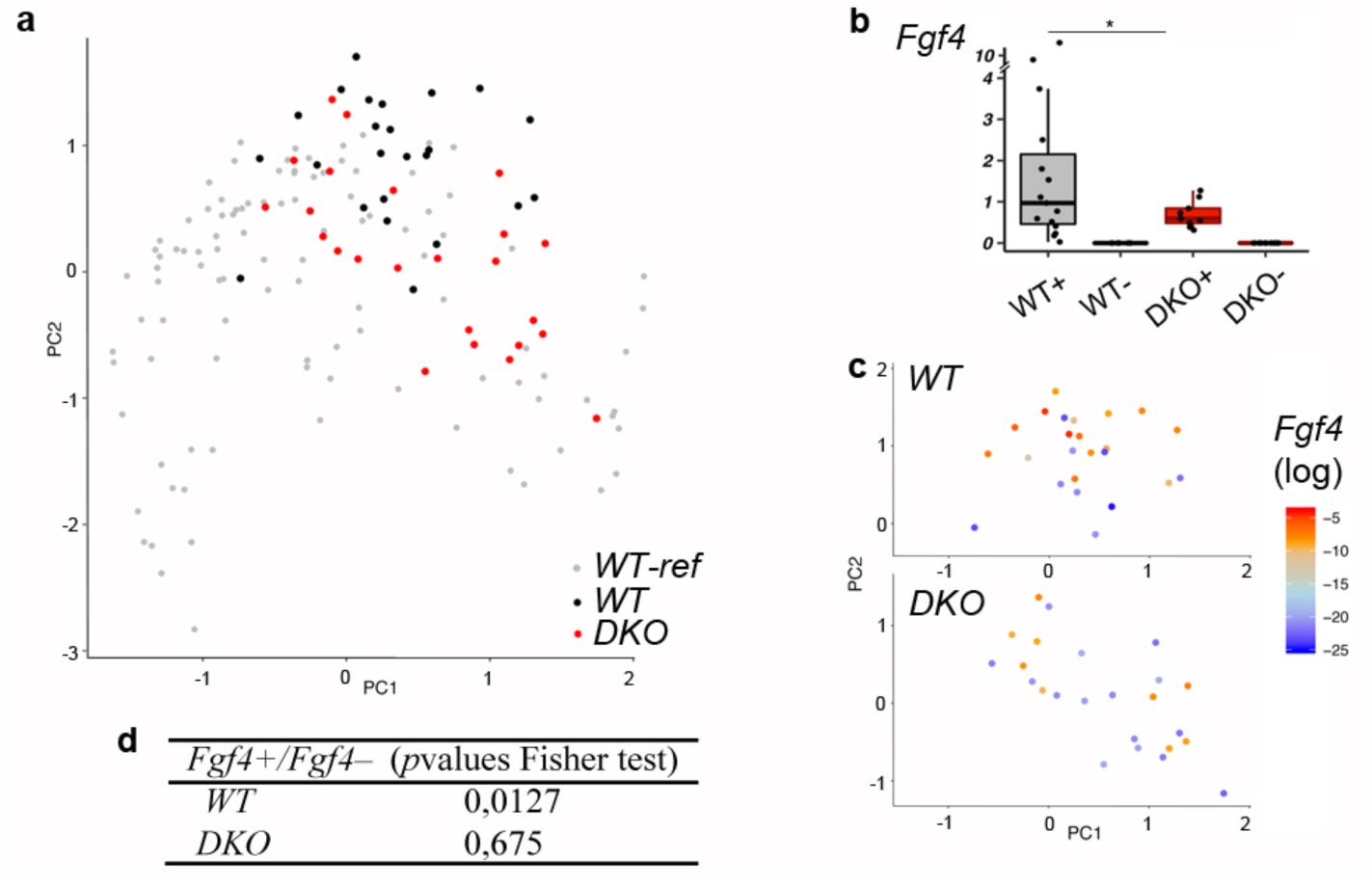
Cell heterogeneity analyses in *DKO* ICMs. **a**, PCA performed with *WT*-Ref (16C, 32C, 64C and 90C), *WT* (32C) and *DKO* (32C) cells (scores: PC1, 20,84%; PC2, 13,13%). **b**, Boxplot showing the single-cell expression levels of *Fgf4* in *WT* and *DKO* cells, subdivided between *Fgf4*+ve and *Fgf4*–ve subpopulations. The asterisk indicates a significant difference between *WT* and *DKO Fgf4*+ve samples distribution (\**p*=0.012; Fligner-Killeen test for homogeneity of variance). **c**, PCA map as shown in (**a**) with *WT* (top panel) and *DKO* (bottom panel) cells only, and with graded colours indicating the level of *Fgf4* expression. **d**, Fisher’s exact test for *Fgf4*+ve and *Fgf4*–ve cells distribution (**c**) after repeated k-means clustering (k=2).

We then carried out single-cell RNA analysis on single and double mutant embryos at stage 32C (SI 4), as our data in the *WT* indicated that this is the onset of *Fgf4+*ve cells and of Epi/PrE differentiation. PCA showed that the cells from *WT* and *DKO* genotypes aligned similarly along the PC1 axis (Fig. 3a; see also Supp data Fig. 4 for PCA analyses on the 4 genotypes) indicating that the *DKO* cells develop at the same pace. Similar to the ICM analysis (SI 1), there were many genes expressed at lower levels in *DKO* embryos whereas others maintained their expression (SI 4). As in *WT*, a subset of cells expressed *Fgf4* in the *DKO* embryos. However, the range of *Fgf4* expression levels in *Fgf4*+ve cells was strongly reduced (Fig. 3b). At stage 32C in the *WT* cells, *Gata6* was similarly expressed in *Fgf4+*ve and *Fgf4*-ve cells (SI 3), suggesting that it probably does not have a differential impact at this stage. Thus, NANOG is required for the high expression of *Fgf4* in individual cells and efficient activation of the pathway, as illustrated by the lack of pERK labelling in *DKO* ICMs (Fig. 1e). The existence of a threshold level for FGF4 activation is supported by data showing that incremental doses of recombinant-FGF4 are required to differentiate PrE cells in *Fgf4* mutants (Kang et al., 2013; Krawchuk et al., 2013). Together, our results indicate that NANOG is necessary to boost *Fgf4* expression in a subpopulation of cells to reach levels required for FGF pathway activation in the neighbouring cells. Thus, the FGF pathway seems to act only downstream of Epi specification.

**Figure 4:**
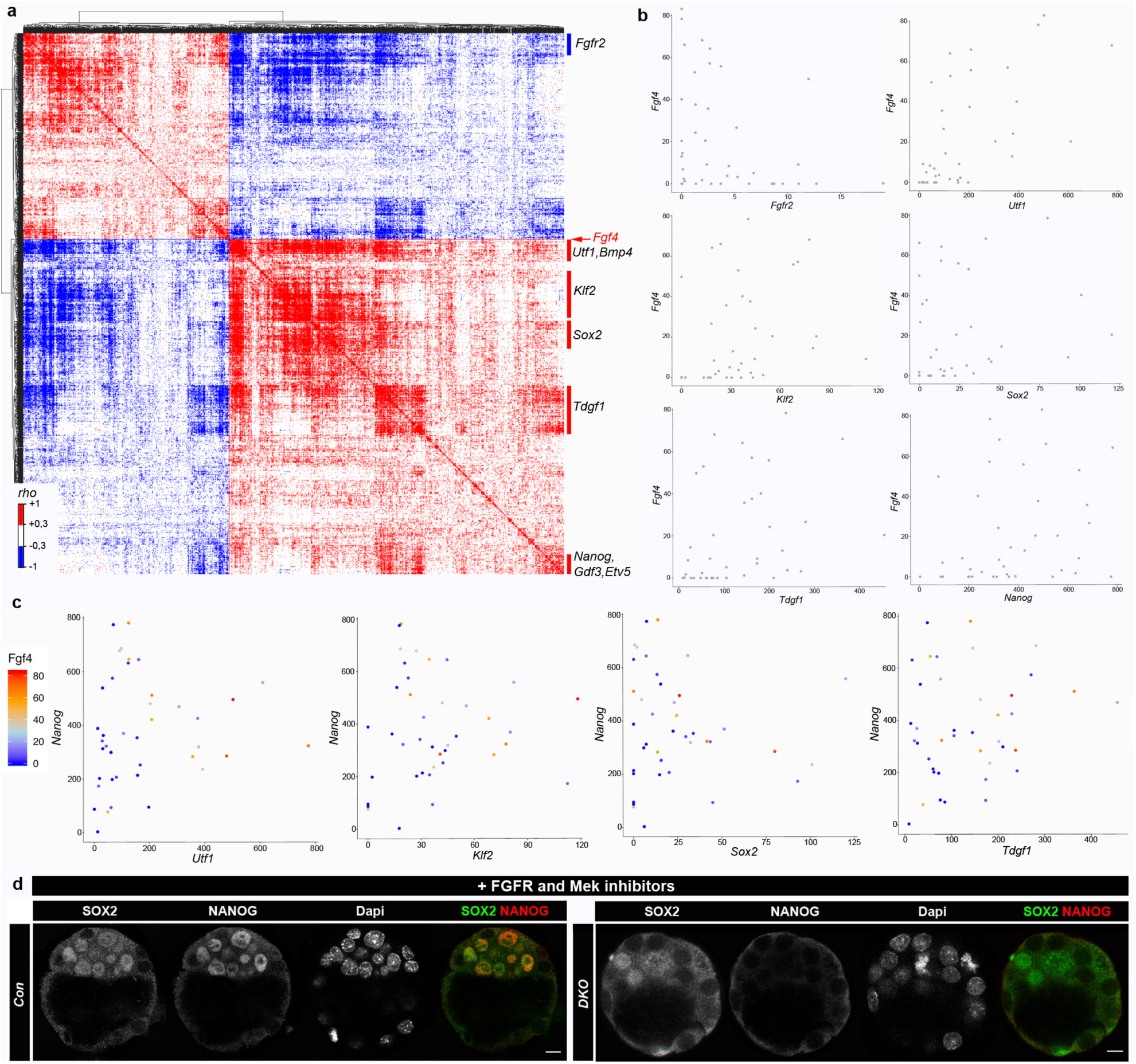
Correlated expression between genes from single cell RNA sequencing. **a**, Spearman correlation matrix for paired expression of the 1434 genes across the 40 ICM cells at stage 32C. Genes are ordered into a hierarchical tree for similarity. Clusters are annotated with representative genes (see SI 6 for detailed map on vectorised PDF). **b**, Quantitative plots comparing the expression of *Fgf4* to representative genes of the clusters shown in (a) in each cell. **c**, Triple quantitative plots comparing in each cell the expression of *Nanog, Fgf4* (through the colour gradient) and an indicated gene. Intensities are expressed in RPKM in **b** and **c. d**, Immunofluorescence labelling for SOX2 and NANOG in *DKO* (right panel, n=6) and control embryos (left panel) cultured with FGFR and MEK1 inhibitors from 8C to 90C stage. Scale bars: 10 µm.

In contrast to the *WT* cells, *Fgf4*+ve cells were not clustered in the PCA of *DKO* cells (Fig. 3c, d) and *Fgf4* expression did not correlate with other genes (Spearman test, *p* >0.05). As a whole, the PCA with *DKO* cells demonstrates that NANOG is required for coordinating the expression of these pluripotency genes to enable the emergence of the Epi state.

Altogether, our results show that correct *Fgf4* expression at the 32C stage requires the presence of NANOG. Yet, at this stage, *Nanog* expression is not strictly restricted to *Fgf4*+ve cells. In addition, *Nanog* is already expressed at the 16C stage with variability between cells (Supp data Fig. 5). Indeed, the NANOG protein can be detected from the 8C stage (Plusa et al., 2008) and is heterogeneously distributed among nuclei around the 16C stage (Dietrich and Hiiragi, 2007). Thus, NANOG alone does not seem to be sufficient to induce *Fgf4* expression, implying that it requires a partner(s). Several genes such as *Zfp42, Tdgf1, Sox21, Pecam1*, had a correlated expression with *Fgf4* at the 32C stage (Supp data Fig. 3c), and maintained their RNA levels in *DKO* ICMs (SI 4). These pluripotency markers showed high heterogeneity in expression levels at 16C (Supp data Fig. 5), however their expression no longer correlated with *Fgf4* at the 16C stage (Spearman test, *p* > 0,05). Thus, if one or several of these genes products are cooperating with NANOG to induce Epi specification, this would be likely through a stochastic activation between the 16C and 32C stages.

Single-cell RNA sequencing analyses have reported in depth cell-to-cell gene expression variation during preimplantation (Boroviak et al., 2015; Deng et al., 2014; Goolam et al., 2016; Mohammed et al., 2017; Posfai et al., 2017; Tang et al., 2011). To further explore possible mechanisms of Epi induction we mined single-cell RNA-seq data, using ICM annotated cells from 32C stage embryos (Posfai et al., 2017). Genes whose expression was correlated with *Fgf4*’s at the 32C stage (SI 5) were selected from the whole transcriptome (≈15K genes), resulting in 888 correlated genes and 546 with an inverse correlation (Spearman test, *p*<0.05). Gene-to-gene expression correlations across the cells were then analysed to identify regulation networks (Fig. 4a, SI 6). The 1434 *Fgf4*-correlating genes were hierarchically clustered into different subgroups, highlighted by *Nanog, Etv5, Utf1, Bmp4, Klf2* and *Tdgf1*, revealing potential co-regulatory mechanisms. Several of these genes were already identified as Epi markers at stage 32C with the Biomark Fluidigm experiments (SI 3), supporting our data. Notably, cells expressing high levels of *Utf1, Klf2* and *Tdgf1*, each representing different subgroups, not only co-expressed *Fgf4* (Fig 4b) but also expressed high levels of *Nanog* transcripts (Fig 4c). A cluster comprising *Fgfr2, Dab2* and *Lrpap1*, which are known PrE markers(Gerbe et al., 2008; Guo et al., 2010; Kurimoto et al., 2006; Ohnishi et al., 2014), showed a negative correlation with *Fgf4* (Fig. 4 a,b). Altogether, these RNAseq data confirmed that already at the 32C stage, a subpopulation of cells has started to acquire an Epi identity. These cells share a gene expression signature determining the earliest Epi state.

These results also revealed candidate regulatory factors for NANOG-mediated induction of the Epi state. In order to identify potential genes that could be involved in the emergence of the Epi lineage from an earlier stage, we carried out an expression correlation analysis on cells at the 16C stage (Supp data Fig. 6b, SI 5, SI 7), when the first ICM cells are produced, using the same dataset (Posfai et al., 2017). At this stage, *Fgf4* transcripts are detected in fewer cells and at lower levels compared to the 32C stage (Supp data Fig. 7a, 7b). Only 68 genes presented a correlated expression with *Fgf4* at both the 16C and 32C stage (SI 5), and the clusters identified at 32C stage were not conserved (Supp data Fig.6b). From the 68 genes, solely *Pou2f3* and *Etv5* code for transcription factors (GO:000368), however their expression at the 16C stage was low compared to the 32C stage (Supp data Fig. 7a, b). Moreover, we could not detect the ETV5 protein before the 32C stage by immunofluorescence (Supp data Fig.8), questioning the functional significance of the low RNA levels at the 16C stage and indicating that this protein is most likely not involved in early ICM cell heterogeneity. Altogether, our single cell RNA analyses revealed several candidate transcription factors for inducing *Fgf4* transcription together with NANOG at 32C, but are either not expressed or randomly expressed at 16C stage.

While our data revealed that levels of *Fgf4* transcripts in individual cells of *DKO* embryos are significantly low, we cannot rule out that their presence still creates some heterogeneity among ICM cells. To exclude this, we cultured litters with *DKO* embryos in the presence of FGFR and MEK inhibitors from the 8C to the 90C stage as previously described (Nichols et al., 2009; Yamanaka et al., 2010). Strikingly, SOX2 (Fig. 4d) as well as SOX21 (Supp data Fig. 9) were still expressed in a mosaic pattern in *DKO*-inhibited ICM cells. This demonstrates that the ICM remains heterogeneous in the absence of *Nanog, Gata6* and the FGF/MEK pathway. Thus, a primary ICM heterogeneity exists before NANOG, GATA6 and the FGF pathway become effective.

Collectively, our results indicate that the emergence of the Epi lineage (*Fgf4*+ve cells) is achieved through the coincidental expression of *Nanog* and other differentially expressed gene(s) in few cells between the 16C and 32C stage. The poor transcription factor expression correlations at stage 16C suggest that these coexpressions appear stochastically. However, NANOG expression variability within ICM cells(Dietrich and Hiiragi, 2007) on its own could be the source of heterogeneity. In that case, NANOG could interact with a homogeneously expressed factor produced from the 16C stage. What initiates NANOG and other factors expression heterogeneity, whether it is due to transcriptional noise or/and to other sources of variability such as cell cycle or post-translational modifications as is proposed for ES cells(Torres-Padilla and Chambers, 2014) is currently unknown in the embryo(Simon et al., 2018). Genes other than transcription factors, such as the ones whose expression correlated with Fgf4’s at the 16C stage could be involved in cell heterogeneity and modulation of NANOG expression or activity. In addition, cell heterogeneity has been found at earlier stages, as expression of proteins including SOX21, CARM1 or PRDM14(Burton et al., 2013; Goolam et al., 2016; Torres-Padilla et al., 2007), and the duration of SOX2 DNA binding(White et al., 2016) are different between blastomeres at the 4C stage and can prefigure inner and outer cells. *Sox2, Sox21, Prdm14* or *Carm1* are not differentially expressed between blastomeres at the 8C stage(Goolam et al., 2016) and their expression is not correlated with *Fgf4*’s at the 16C stage (SI 5). Therefore, if these genes are involved in *Fgf4* expression heterogeneity, there must be a relay mechanism between the 4C and 32C stages, presumably at the posttranscriptional level. *Sox21* and *Prdm14* mutant mice are viable(Kiso et al., 2009, 21; Yamaji et al., 2008) and the number of NANOG-expressing cells is unchanged in *Sox2* maternal-zygotic mutant embryos at 90C(Wicklow et al., 2014), indicating that these genes are probably not required to induce the initial Epi cells. Interestingly, *Carm1* maternal-zygotic mutant blastocysts have a smaller total number of cells and a smaller number of NANOG-expressing cells(Hupalowska et al., 2018). It would thus be interesting to examine how *Fgf4* expression is induced and how Epi and PrE cells are specified in these embryos.

Thus, it remains unclear whether the emergence of Epi cells results from the stochastic co-expression of different factors or from the inheritance of an earlier cell heterogeneity. Nevertheless, the asynchrony of Epi, and thus PrE, cell specification that is taking place between 32C and 90C stages^2,3,38^ can be explained by the variability of gene expression that drives the coincidental expression of NANOG with other transcription factor(s).

We also show that ICM cell heterogeneity is present in the absence of NANOG, GATA6 and the FGF pathway, and that effective FGF signalling is a consequence of Epi specification through NANOG expression. Thus Epi specification is not induced by the absence of FGF signalling, although it is a necessary condition for its occurrence, but by other factors driving early ICM heterogeneity. Thus, a primary cell heterogeneity is present independently of the FGF pathway, and is then emphasized by NANOG, GATA6 and FGF activities to translate this early cell heterogeneity into cell differentiation. Human and other mammalian embryos are less sensitive than rodents to the FGF pathway^47–49^ for establishing the “salt and pepper” expression of GATA6 and NANOG expression. Thus, this mechanism of primary ICM heterogeneity that we propose for the emergence of the Epi cells induced by NANOG and coincidental expression of currently unknown partner(s) independently of the FGF pathway, could be conserved amongst mammals.

## Methods

### Embryos collection and staging

Experiments were performed in accordance with French and EU guidelines for the care and use of laboratory animals. Animals were housed in a pathogen-free facility under a 12-hour light cycle. All embryos used in this study were produced from natural mattings.

Embryos were collected at 8-cell stage (8C), 16C, 32C, 64C and 90C (corresponding to 2.5, 3.0, 3.25, 3.5, and 3.75 days of embryonic development respectively). For the single cell analysis on *WT* embryos (*WT*-Ref) from the CD1 strain (Charles River or Janvier), embryos were also staged using littermates total cell counts.

### Genotyping

Single and double mutant embryos were obtained from heterozygous intercrosses between *G*^*-/+*^ (*Gata6*^*tm2.1Sad*^)(Bessonnard et al., 2014; Sodhi et al., 2006) and *N*^*-/+*^ (*Nanog*^*tm1Yam*^)(Mitsui et al., 2003) lines (backcrossed to CD1 mice for more than 10 generations). In a subset of embryo cultures, *Gata6*^*tm2.1Sad*^ allele with the *Zp3-Cre* transgene (de Vries et al., 2000) were used to increase the number of double mutants. Mendelian ratios were obtained for the different genotypes.

Mice and embryos were genotyped as previously described (Bessonnard et al., 2014; Chazaud and Rossant, 2006; Frankenberg et al., 2011). Primer pairs for *Gata6* genotyping (Sodhi et al., 2006), using the Goldstar Taq polymerase (Eurogentec) were: 5’-AGTCTCCCTGTCATTCTTCCTGCTC-3’associated to5’-TGATCAAACCTGGGTCTACACTCCTA-3’ for the mutated allele and 5’-GTGGTTGTAAGGCGGTTTGT-3’ associated to 5’–ACGCGAGCTCCAGAAAAAGT–3’ for the wild-type allele. Primer pairs for *Nanog* genotyping (Mitsui et al., 2003), using the Taq polymerase (Invitrogen or Promega) were: 5’-CAGAATGCAGACAGGTCTACAGCCCG-3’ coupled with either 5’-AATGGGCTGACCGCTTCCTCGTGCTT-3’ for the mutated allele or 5’-GGCCCAGCTGTGTGCACTCAA-3’ for the wild-type allele.

### Embryo cultures and immunostaining

Embryos were flushed with M2 (Sigma Aldrich) and cultured in KSOM medium (Millipore) for the indicated time points. Embryos were treated with FGFR inhibitor (PD173074, Sigma Aldrich) at 100nM and MEK1 inhibitor (PD0325901, Sigma Aldrich) at 500nM (Frankenberg et al., 2011). Blastocysts immunostainings were performed as previously described (Chazaud et al., 2006) (see antibodies list, Supp data Table 1)

Images were captured with confocal microscopes Leica SPE, SP5 and Sp8 using 40X objectives. Images were analysed with ImageJ (NIH) and Imaris (Bitplane) softwares. Cell counts were performed as previously described (Bessonnard et al., 2014). ICM cell numbers were counted by subtracting the number of CDX2 expressing cells to the total (dapi-labelled) cell number. Unless indicated, images show single confocal image z-slices.

### RT-qPCR on single ICM

RT-qPCR on single ICM was adapted from described protocols (Ralston et al., 2010; Tang et al., 2010).

### Boxplots presentation

The edges of the box represent 25th and 75th quartiles. The median is represented by the central line. The whiskers extend to 1.5 times the interquartile range (25th to 75th percentile) Cells are plotted individually in single-cell experiments.

### Single cell isolation and RT-qPCR/Fluidigm analysis

Isolated ICM cells (after immunosurgery) and morulae (16C) were incubated 10 minutes in 1X TrypLE™ Express Enzyme (Gibco) at 37°C and cells were isolated by repeated mouth pipetting using pulled capillaries of serially smaller diameter openings. Each single cell was collected in 5µl of 2X Reaction Mix (Invitrogen, CellsDirect One-Step qRT-PCR Kit) and stored at −80°C or processed immediately.

Single-cell qPCR on a Fluidigm Biomark system (GENTYANE facility) was carried out according to the manufacturer’s instructions. Cells with absent or low Ct values for housekeeping genes were removed from analysis (5%). Ct values were normalized using the 2-ΔCt method. In boxplots, values are relative to the mean of all WT 32C cells (*WT*-ref and *WT* from the transgenic background when available) to be able to compare between genotypes and stages. Cells were considered in *Fgf4*–ve subpopulations when CT values were above 35 and all other cells were placed in *Fgf4*+ve subpopulations.

### Statistical analyses

Statistical tests were performed with Graphpad software or R. Statistical significance was assessed using the Wilcoxon-Mann-Whitney test (non parametric) on expression levels (see the different Excel sheets in SI 8). A Fligner-Killeen non-parametric test for homogeneity of variance was performed to analyse the distribution of *Fgf4*+ve cells between *WT* and *DKO* cells.

### Generation of heatmaps

To construct heatmaps, we used R with the package « pheatmap » (Kolde R. 2015. Package ‘pheatmap’: https://CRAN.R-project.org/package=pheatmap). We made a log base 2 transformation on cells expression values. Then we applied a hierarchical clustering algorithm on both cells and genes values to obtain the final heatmap with dendogram showing distances.

### PCA analyses

Principal component analyses were performed using R package « pcaMethods » using « bpca » method (Stacklies et al., 2007) to compute component scores on log2 expression values from stages and genotypes. Scatter plots showing the distribution of cells and of genes between the two main components (PC1 and PC2) were produced.

To analyse the clustering between *Fgf4+*ve and *Fgf4*-ve cells, we first used a repeated k-means clustering method with k=2 on each sample (32C, 64C and 90C *WT*-Ref, *WT* and *DKO*). We then analysed the distribution of *Fgf4+*ve and *Fgf4*-ve cells within the clusters using a Fisher’s exact test to examine the significance of the association between the two kinds of classification.

### Single-cell RNA-seq expression correlation analyses

Single-cell expression data were extracted from (Posfai et al., 2017), taking 40 ICM cells from the 32C stage (early and late) and 33 inner cells from the 16C stage (already defined by Posfai and collaborators)(Posfai et al., 2017).

All correlation *rho* scores and *p* values were performed using R package “HMisc” with the Spearman method. To obtain correlation heatmap for 32C stage, we first computed all expression correlations between *Fgf4* and all the genes at 32C stage. Genes that had *rho* score >= 0,3 (corresponding to *p* <= 0,05 for 40 samples) were kept for building the matrix.

Correlation scores in between these genes were computed and discretized in 3 individuals’ values: 1= correlated (*rho* score >= 0,3), −1 = anticorrelated (*rho* score <=−0,3), 0 = no correlation (otherwise).

To obtain correlation matrix for 16 cells stage we used the 1434 genes selected as correlated or anticorrelated to *Fgf4* expression at 32C stage. All correlation scores between these genes were computed and correlation was considered significant when *rho* score>= 0,345 (corresponding to *p* <= 0,05 for 33 samples). Then we discretized these values in 3 individuals’ values: 1= correlated (*rho* score >= 0,345), −1 = anticorrelated (rscore <=-0,345), 0 = no correlation (otherwise).

Heatmaps were generated using R package “pheatmap”.

### Comparison Gene plot

Comparison gene plots were made using R package “ggplot2” using “geompoint()” function.

## Acknowledgements

Images were acquired and treated at the CLIC imaging facility (Clermont-Fd). C.C. thanks C. Pléver for genotyping the mice. Authors want to thank V. Mirouse, B. Cox, T. Hiiragi and J. Rossant for insightful comments on the manuscripts. Authors acknowledge Life Science Editors for editing assistance.

## Author contributions

CC, NA and SC conceived and designed the experiments. CC, NA, SC, CD, LVE performed the experiments. YR performed the bioinformatic analyses. PP designed semi-automated image analyses. CC, MCT, NA, YR, CD and SC analysed the data. CC, NA and MCT wrote the manuscript. C.C. was supported by the Agence Nationale de la Recherche (ANR-14CE11-0017 PrEpiSpec) and the FRM. MCT was supported by was supported by the Institut Pasteur and the Agence Nationale de la Recherche (ANR-10-LABX-73-01 REVIVE and ANR-14CE11-0017 PrEpiSpec).

## Supplemental data

**Supp data Figure 1:**
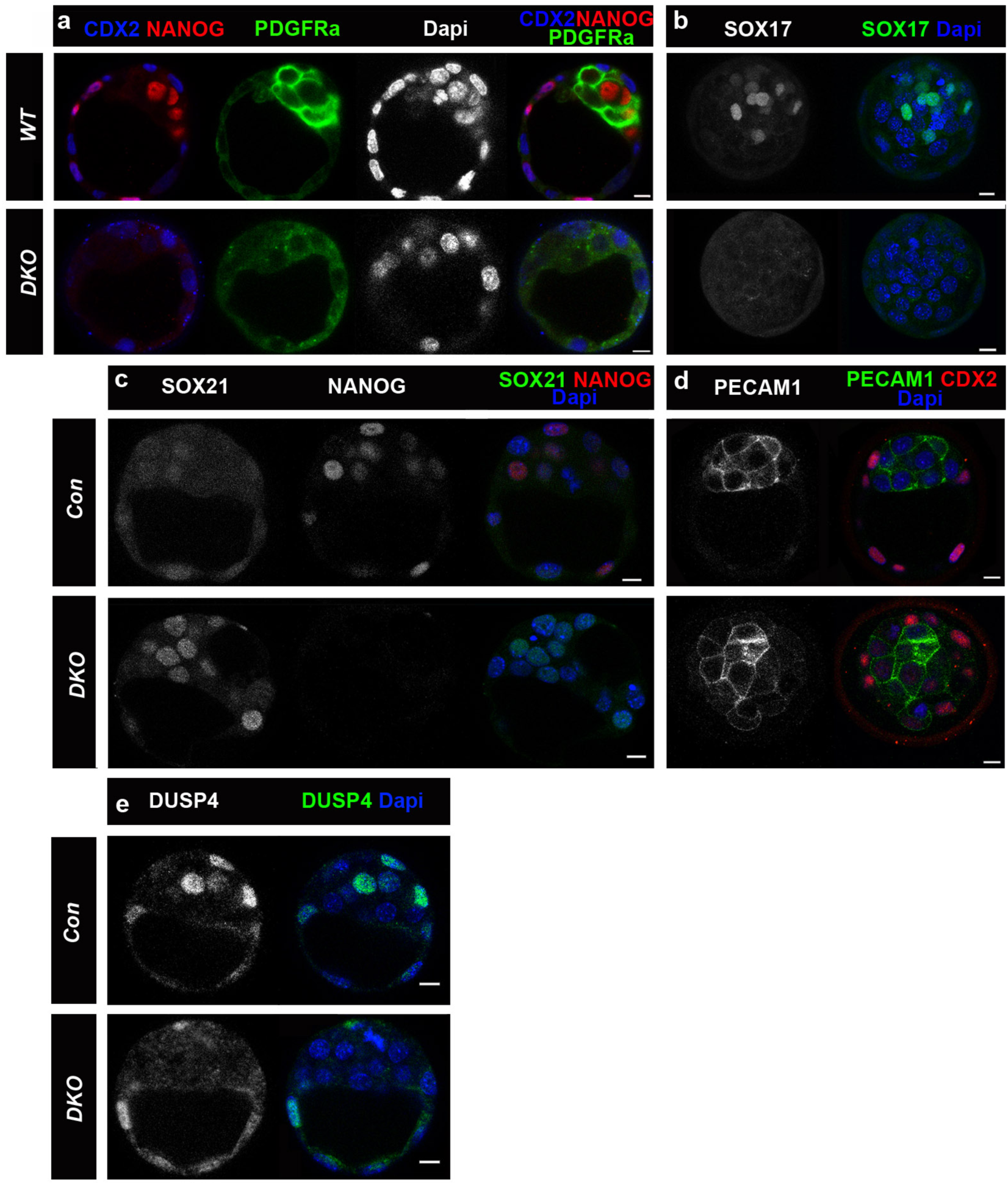
Immunolocalisation of different markers in *DKO* blastocysts. **a**, CDX2, PDGFRa and NANOG expression at 90C stage. **b**, Projected confocal slices showing the PrE marker SOX17 expression in representative control and its absence in *DKO* embryos at 64-90C stages. **c, d**, The pluripotency markers PECAM1 and SOX21 are expressed in both control and *DKO* embryos at 64-90C stages. Note that SOX21 expression is stronger in *DKO* embryos as its expression decays around 90C in *WT* embryos. **e**, DUSP4 expression is down regulated in *DKO* ICMs at 64-90C stages. DUSP4 can be detected in some TE cells but not ICM cells. Scale bars: 10 µm. Number of *DKO* embryos analysed (fully penetrant phenotypes): TE (CDX2) n>8; PrE (SOX17 and PDGFRa) n=3; SOX21 n=4; PECAM1 n=2 and DUSP4 n=3

**Supp data Figure 2:**
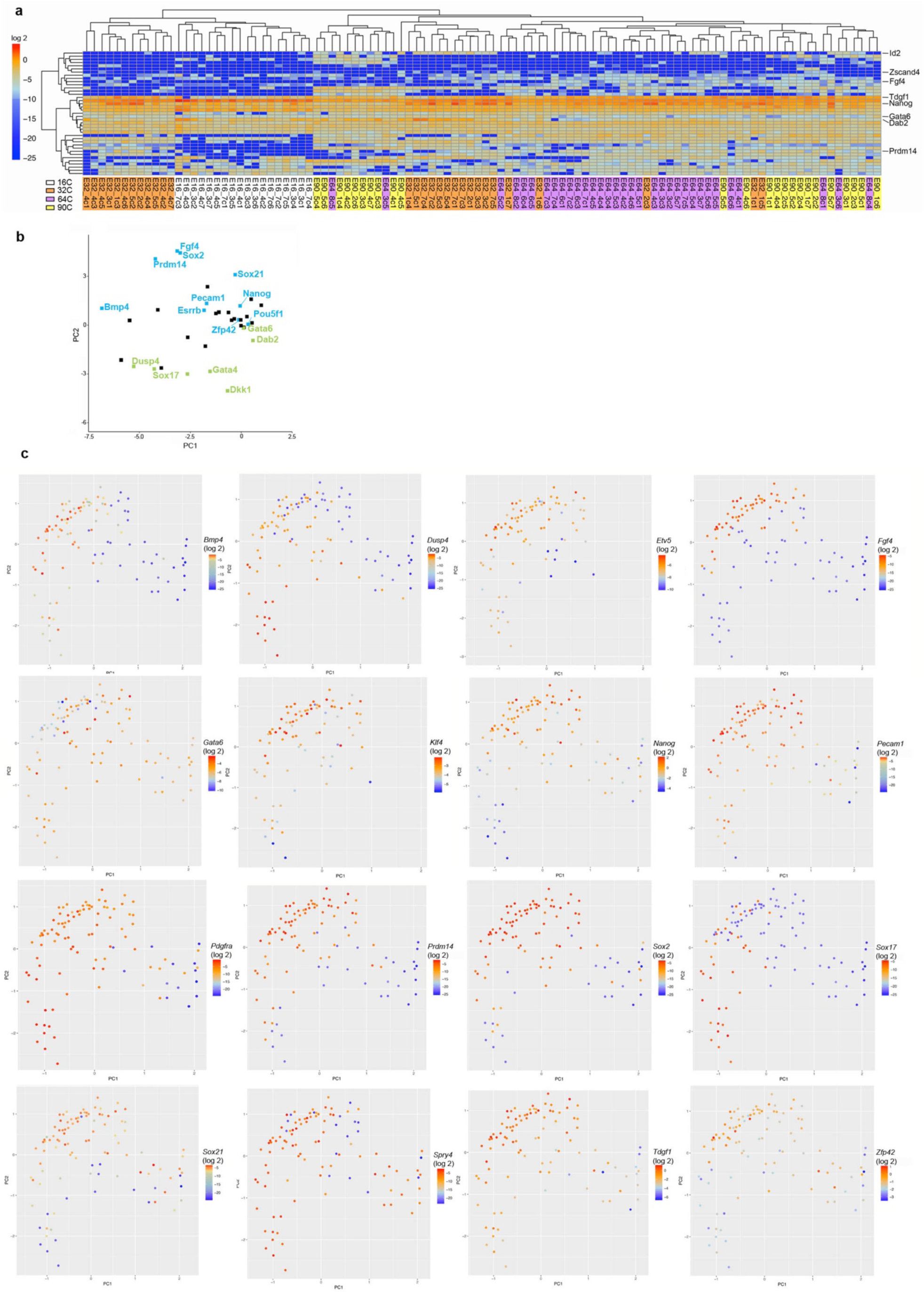
**a**, Heatmap indicating the relative expression levels of 44 genes in WT single ICM cells at the four stages, and ordered through an unsupervised hierarchical clustering. **b**, PCA loading map (see Fig 2c) showing the distribution of the measured gene expression levels. Epi and PrE genes are labelled in blue and green respectively. **c**, PCA map shown in (Fig. 2**c**) with graded colours indicating the level of different genes expression in each cell at the four stages. For *Etv5* and *Klf4*, experiments on 16C stage cells were not carried out.

**Supp data Figure 3:**
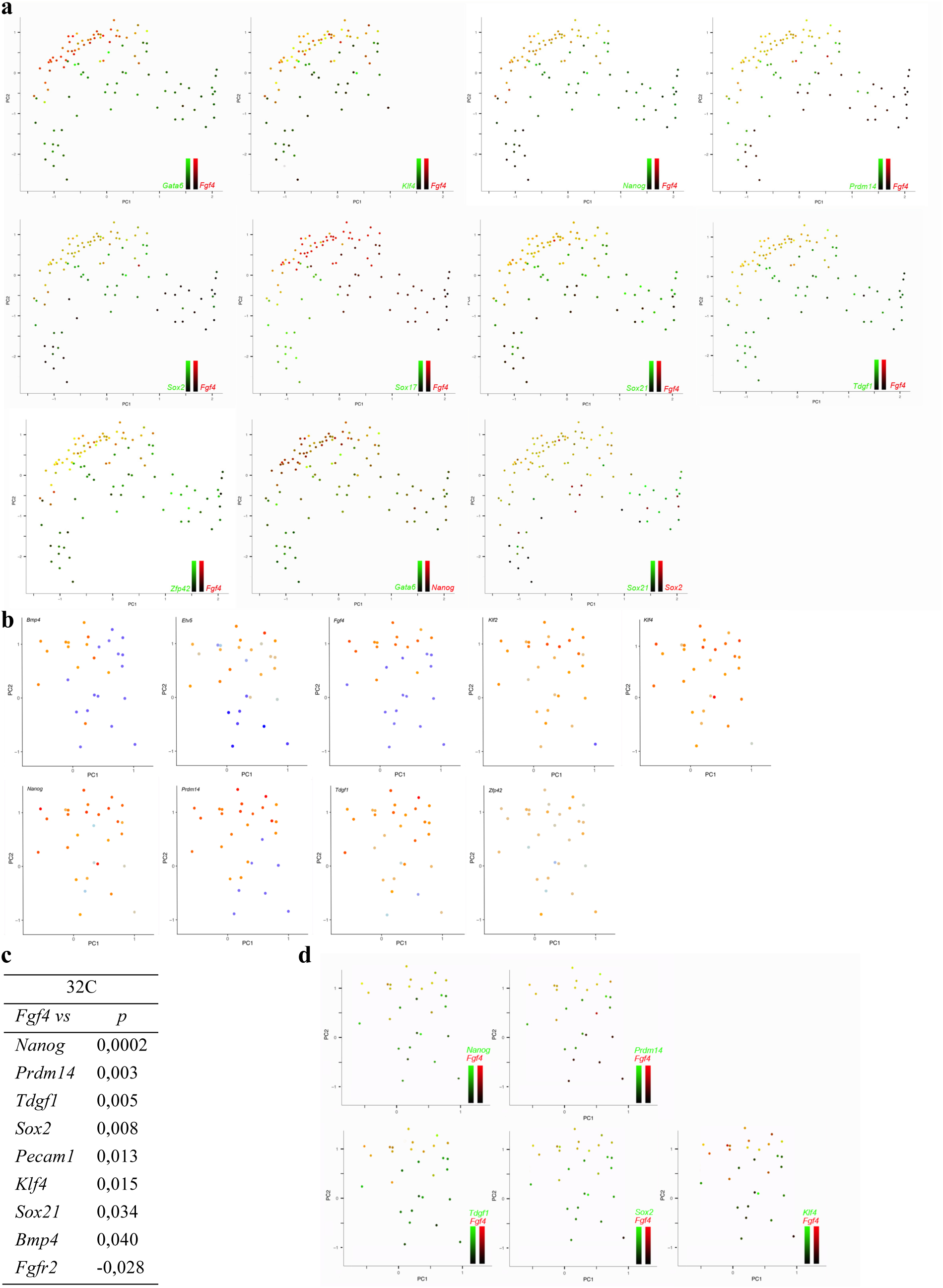
Co-expression of genes in individual cells. **a**, Paired genes expression in red and green to observe their colocalisation in yellow on the PCA map. For each gene, intensity scales (log2) are shown in Supp data figure 3d. **b**, PCA map focused on the 32C stage with different genes expression gradient (see ext data Fig. 2c). **c**, Positive or negative correlation (Spearman test) of *Fgf4* versus genes expression in 32C stage cells (results with significant values are shown). **d**, examples of paired genes expression at 32C stage as in **a**.

**Supp data Figure 4:**
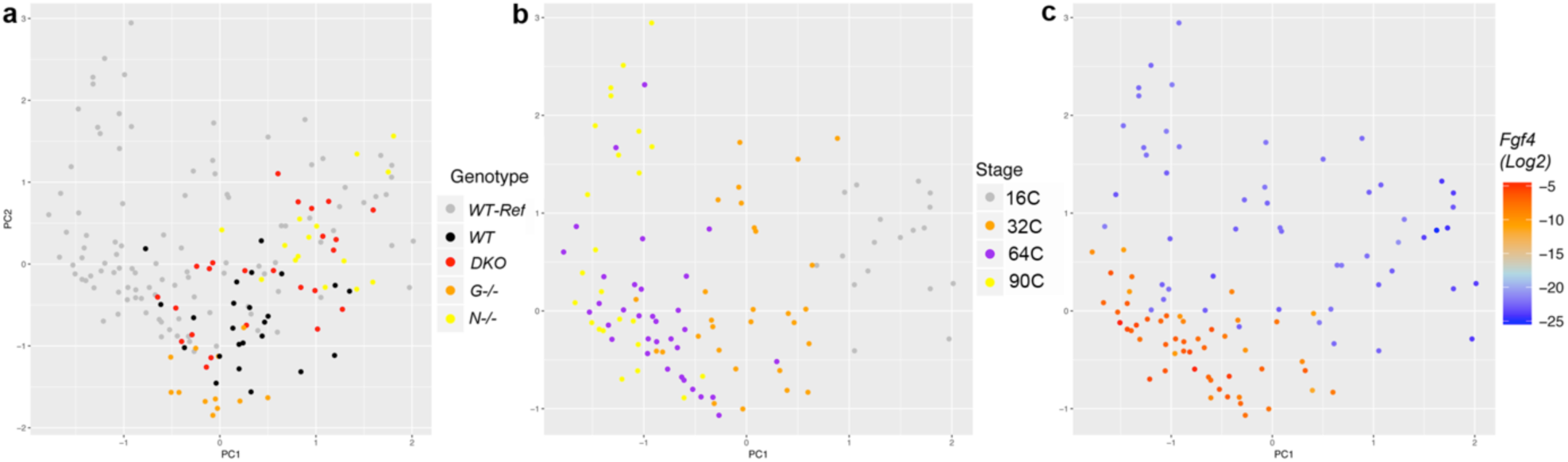
**a**, PCA maps built with the cells of the four genotypes (*WT, N*^-/-^, *G*^-/-^ and *DKO*) as well as with cells of *WT-Ref*, used as reference. Scores: PC1: 19,38% PC2: 13,13%. **b**, same PCA map, showing only *WT-Ref* cells according to the stages. **c**, same PCA map, showing only *WT-Ref* cells with *Fgf4* expression intensities.

**Supp data Figure 5:**
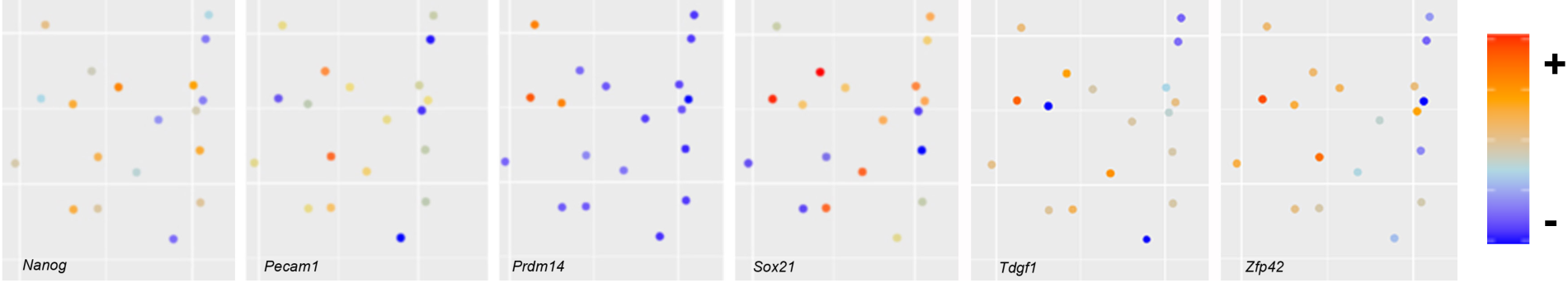
PCA map of Fig. 2g showing only the 16C stage with different genes expression gradient (as in Supp data Fig. 2c). *Fgf4* is not detected at this stage (see Supp data Fig.2c).

**Supp data Figure 6:**
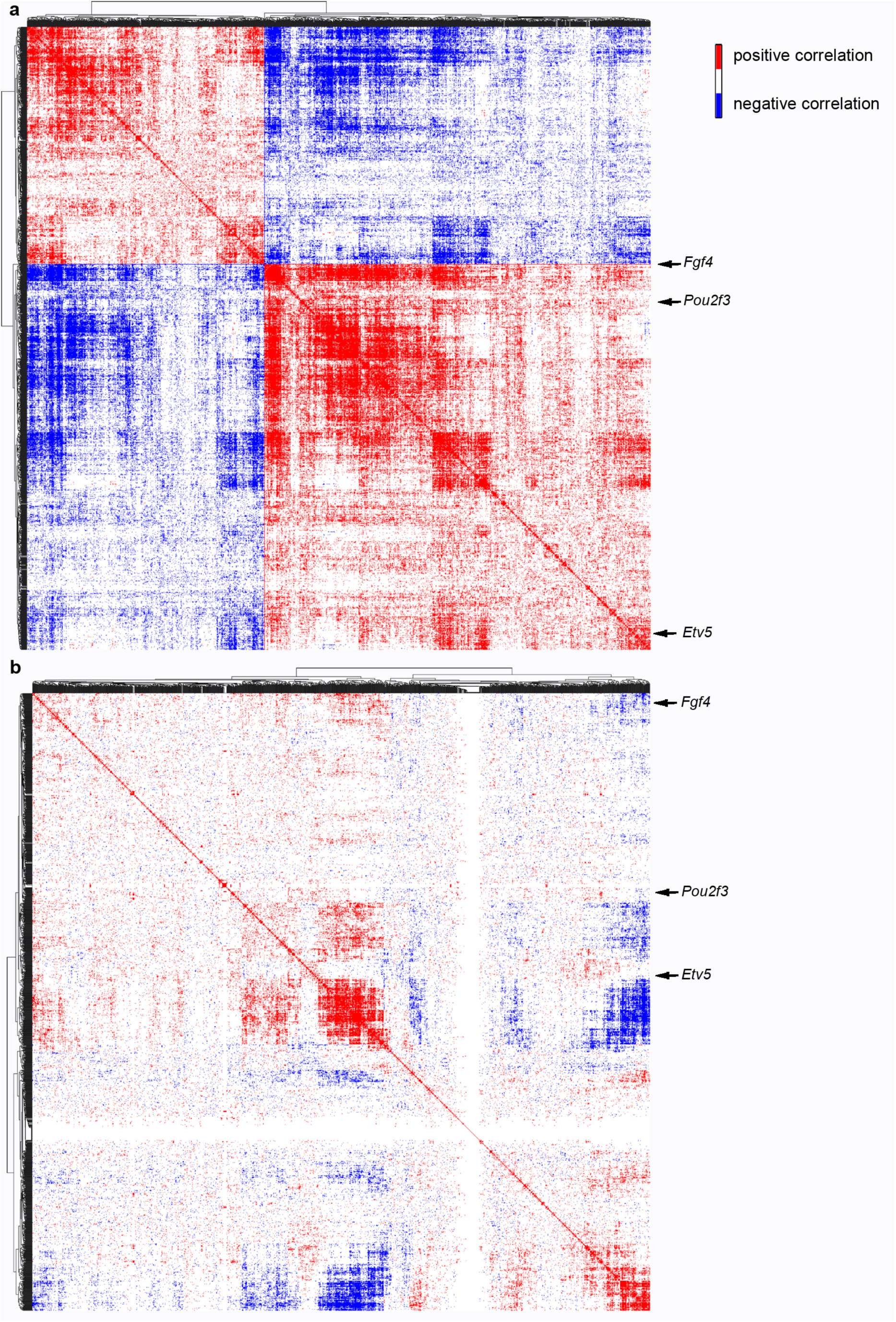
Spearman correlation matrix for gene expression from RNAseq data. 1434 out of 15 713 genes were selected through their correlated expression to *Fgf4*’s (Spearman test, *p* < 0,05, see SI 6 for exact values). Expression correlation heatmaps were built with all pairs of the selected genes from 40 ICM cells at the 32C stage (**a**) and 33 inner cells at the 16C stage (**b**) (see SI 6 and SI 7 for detailed maps on vectorised PDFs). Genes are ordered into a hierarchical tree for similarity. Spearman correlation tests (*p* < 0,05).

**Supp data Figure 7:**
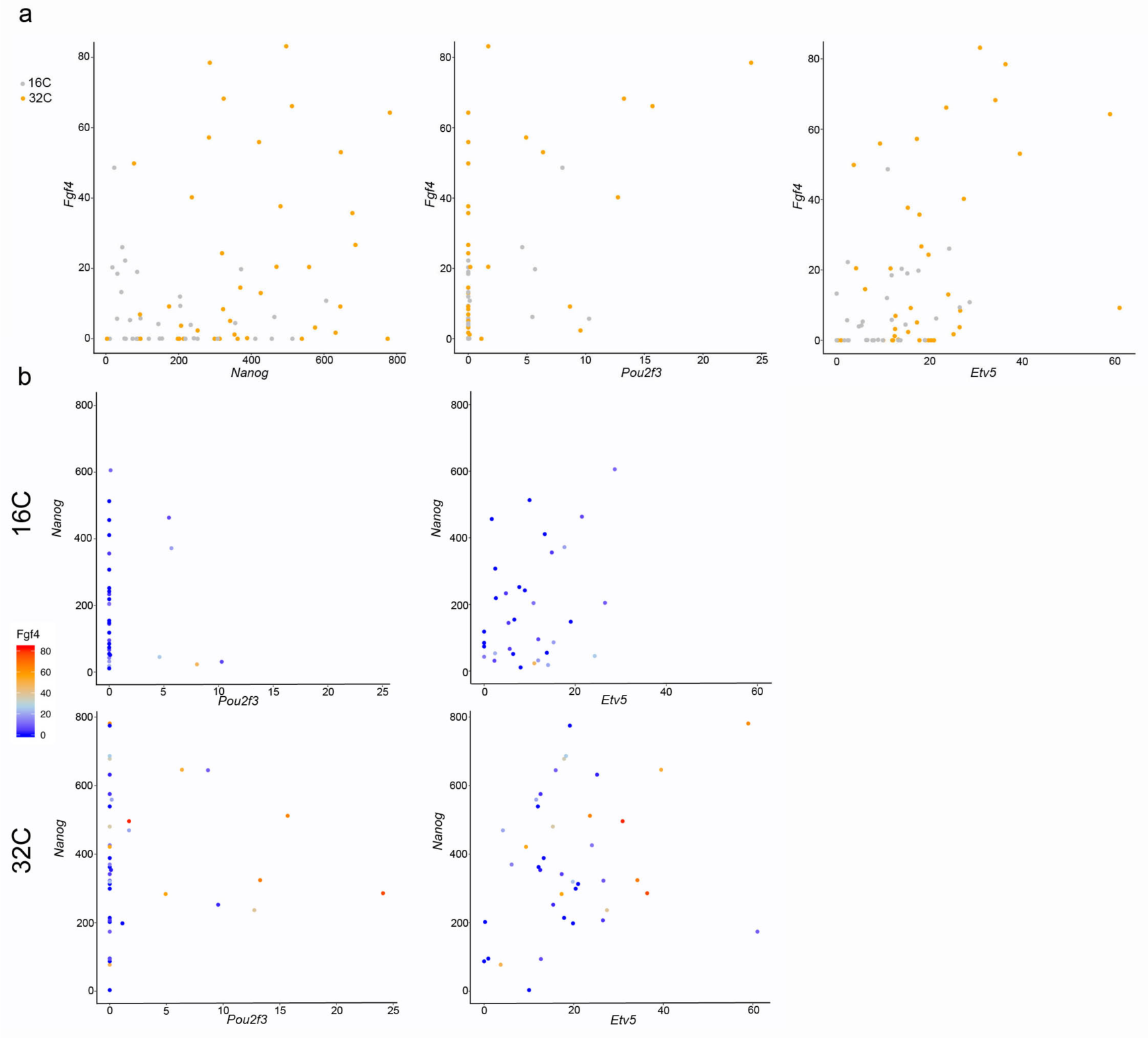
(**a**) Quantitative plots comparing the expression of *Fgf4* to *Nanog, Pou2f3* and *Etv5*’s in individual cells at 16C and 32C stages. Intensities are expressed in RPKM. Very few cells are expressing *Pou2f3* at the 16C stage while *Etv5* and *Fgf4* are mutually expressed in the same cells at both stages, also illustrated by their correlation coefficient (*p*= 0,01 at 16C and 32C stages). (**b**) Triple quantitative plots comparing in each cell the expression of *Nanog, Fgf4* (through the colour gradient) with *Pou2f3* or *Etv5* at the 16C (Top panels) or 32C (bottom panels) stages. Intensities are expressed in RPKM.

**Supp data Figure 8:**
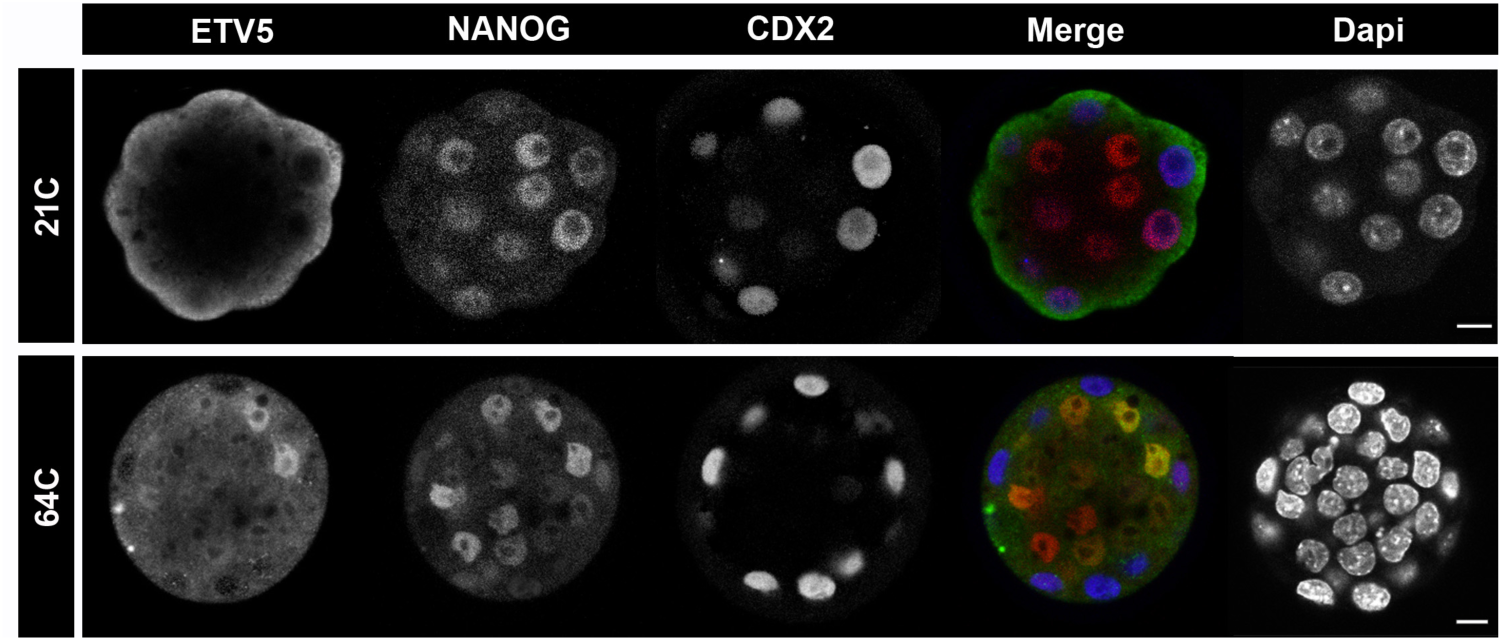
Expression of ETV5, NANOG and CDX2 just before cavitation (21C, top panel) and at the 64C stage (bottom panel, section throughout the ICM). ETV5 cannot be detected before stage 32C, there is a weak expression in few ICM cells in few embryos at stage 32C (not shown) and from the 64C stage expression can be seen in some ICM cells, mainly colocalized with NANOG. scale bar: 10 µm.

**Supp data Figure 9:**
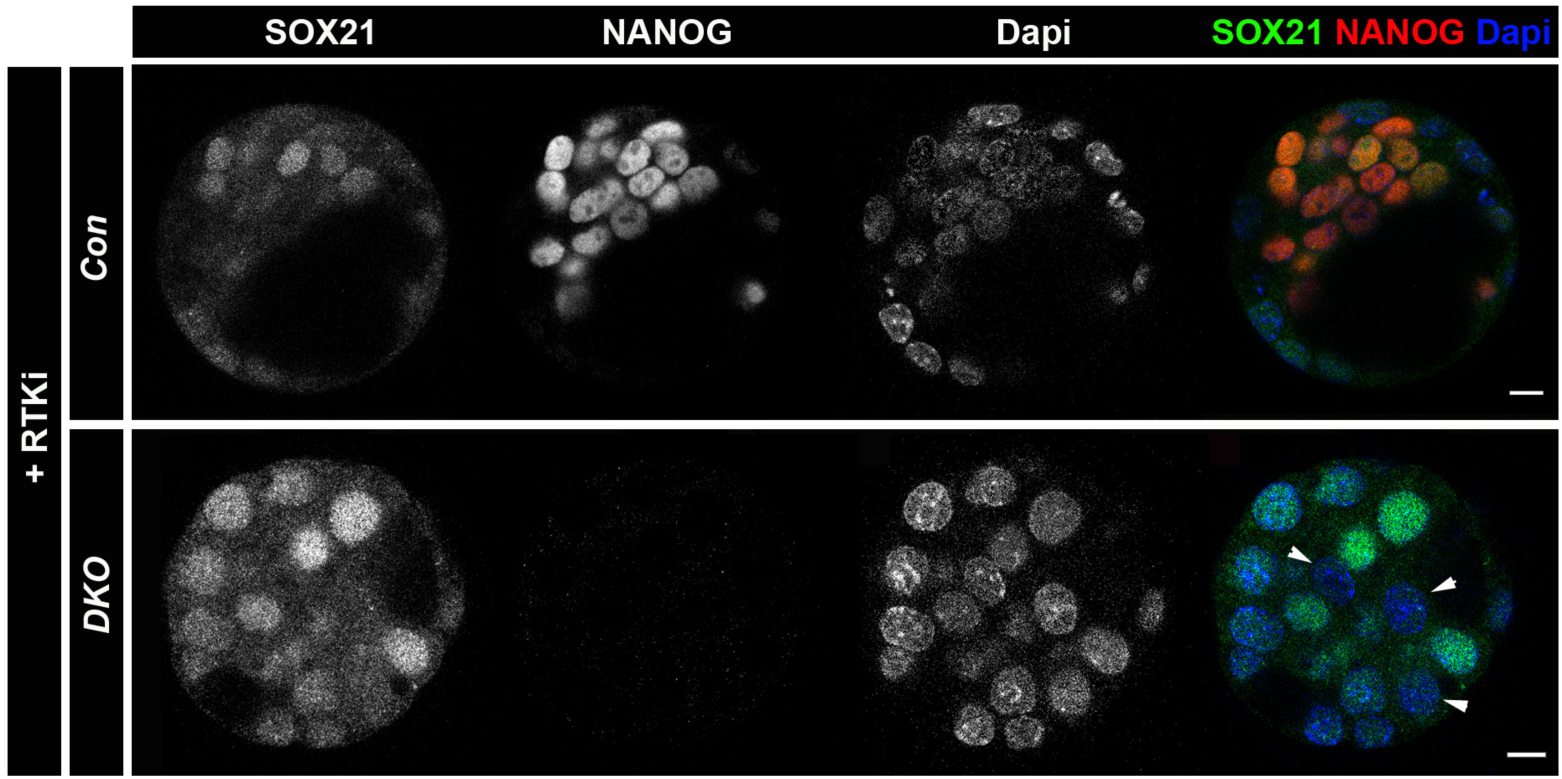
*DKO* and control embryos cultured with FGFR and MEK1 inhibitors from 8C to 90C stage. SOX21 remains expressed heterogeneously in absence of *Nanog, Gata6* and the FGF pathway (n=3). Arrowheads point toward unlabelled SOX21 ICM nuclei. Scale bars: 10 µm.

